# Histone H3 tail modifications required for meiosis in *Saccharomyces cerevisiae*

**DOI:** 10.1101/2024.12.09.627563

**Authors:** Amy Prichard, Marnie Johansson, David T. Kirkpatrick, Duncan J. Clarke

**Affiliations:** Department of Genetics, Cell Biology & Development, University of Minnesota, Minneapolis, MN, USA

**Keywords:** Histone H3, Meiosis, phosphorylation, H3T3, sporulation, histone modifications

## Abstract

Histone tail phosphorylation has diverse effects on a myriad of cellular processes, including cell division, and is highly conserved throughout eukaryotes. Histone H3 phosphorylation at threonine 3 (H3T3) during mitosis occurs at the inner centromeres and is required for proper biorientation of chromosomes on the mitotic spindle. While H3T3 is also phosphorylated during meiosis, a possible role for this modification has not been tested. Here, we asked if H3T3 phosphorylation (H3T3ph) is important for meiotic division by quantifying sporulation efficiency and spore viability in *Saccharomyces cerevisiae* mutants with a T3A amino acid substitution. The T3A substitution resulted in greatly reduced sporulation efficiency and reduced spore viability. Analysis of two other H3 tail mutants, K4A and S10A, revealed different effects on sporulation efficiency and spore viability compared to the T3A mutant, suggesting that these phenotypes are due to failures in distinct functions. To determine if the spindle checkpoint promotes spore viability of the T3A mutant, the *MAD2* gene required for the spindle assembly checkpoint was deleted to abolish spindle assembly checkpoint function. This resulted in a severe reduction in spore viability following meiosis. Altogether, the data reveal a critical function for histone H3 threonine 3 that requires monitoring by the spindle checkpoint to ensure successful completion of meiosis.

## Introduction

Meiosis generates haploid progeny from diploid cells from two rounds of chromosome segregation and is highly conserved throughout eukaryotes, from yeast to humans and many other organisms [1]. In meiosis I, cells segregate the homologous chromosomes, and in meiosis II the sister chromatids are segregated. In *Saccharomyces cerevisiae*, meiosis is accompanied by sporulation which reorganizes the mother cell cytoplasm to produce individual plasma membranes and cell walls to encapsulate the four haploid cells (spores) [2]. The nature of this process provides the opportunity to examine the ability of cells to complete meiosis successfully, producing mature spore asci, and allows examination of recombination frequencies and chromosome segregation fidelity.

Histones are the central proteins that make up nucleosomes. DNA is wrapped around nucleosomes, allowing the chromatin to take on higher structure. Each nucleosome is an octamer that contains two heterodimers of histone H2A and histone H2B and a heterotetramer of two histone H3 and two histone H4 proteins [3]. Histones are globular proteins with largely unstructured N-terminal tails [4]. The tails are approximately 25-35 amino acids in length with a high proportion of basic residues [3]. Both the N-terminal tails and the globular histone core domains are subject to a number of post-translational modifications [5]. Commonly, histones can be phosphorylated, methylated, and acetylated on their N-terminal tails. Modifications to the histone tails affect how different proteins can access the DNA [5].

A conserved feature of meiosis in eukaryotes is that it is accompanied by histone post-translational modifications and accumulating evidence indicates that these modified residues orchestrate meiosis-specific gene expression patterns, chromosomal structural changes and recombination, as well as kinetochore-microtubule attachment ahead of meiotic chromosome segregation [6–11]. Histone modifications can have numerous effects such as changing chromatin structure and recruiting other proteins to the chromatin, which in turn controls cellular processes including transcription, DNA repair, and cell division [12, 13]. During mitosis histone H3 is phosphorylated at threonine 3 (T3) by Haspin kinase [14]. This modification primarily occurs on histone H3 proteins at the inner centromeres of chromosomes [15–18] where it serves to recruit the Chromosome Passenger Complex (CPC) including Aurora B kinase [17, 19–22]. Given the essential roles of Aurora B at centromeres in mitosis, Haspin-mediated T3 phosphorylation is needed for proper chromosome alignment on the spindle and accurate chromosome segregation [15, 16].

H3T3 is phosphorylated at centromeres during meiosis I and II in mouse oocytes [23]. Although a possible requirement for T3 phosphorylation in meiosis has not been directly tested, inhibition of Haspin kinase causes defects in kinetochore-microtubule attachment and aneuploidy is observed following the completion of meiosis [23, 24]. Recent studies also detected H3T3 phosphorylation in mitotic *S. cerevisiae* cells [21]. Proper localization of the yeast ortholog of Aurora B kinase (Ipl1) was found to be disrupted when T3 phosphorylation was abrogated during mitosis [21]. However, H3T3 has not been examined for a possible role in meiosis in yeast.

Yeast serves as a powerful model to study meiosis and previous work identified histone H3 and H4 residues required for yeast sporulation [25]. To determine if H3T3ph is required for successful meiosis, we generated yeast mutants where histone H3 T3 is substituted with alanine (T3A) to prevent phosphorylation. To characterize T3A homozygous and heterozygous mutants, we quantified sporulation efficiency, chromosome segregation, and spore viability. In parallel, we examined H3 S10A homozygotes and heterozygotes, and H3 K4A homozygotes and heterozygotes, representing other prominent post-translationally modified histone H3 N-terminal tail residues [11]. This analysis revealed differential roles of these residues in meiosis. We also examined *mad2Δ* mutants that have a non-functional spindle assembly checkpoint [26]. This revealed the spindle assembly checkpoint is crucial for successful meiosis in the H3 T3A mutant.

## Materials and Methods

### Yeast strains

The strains used were all derived from the *Saccharomyces cerevisiae* W303 laboratory strain. The yeast strains that were used are listed in Table 1.

**Table 1.** Yeast strains used in this study. All yeast strains used in this study are derived from W303. The strain numbers are listed with the mating type (MAT), experimental group, and strain description (genotype).

| Number | MAT | Group | Genotype |
| --- | --- | --- | --- |
| DCY2459 | a | Wild type | *ade2-1, ura3, leu2, his3, trp1* |
| DCY 2460 | $\alpha$ | Wild type | *ade2-1, ura3, leu2, his3, trp1* |
| DCY 4595 | $\alpha$ | T3A | *hht1-hhf1::HYG<sup>R</sup>, hht2-T3A-HHF2(URA3), ade<sup>-</sup>, leu2<sup>-</sup>, his3<sup>-</sup>* |
| DCY 4596 | a | T3A | *hht1-hhf1::HYG<sup>R</sup>, hht2-T3A-HHF2(URA3), ade<sup>-</sup>, leu2<sup>-</sup>, his3<sup>-</sup>* |
| DCY 4706 | a/$\alpha$ | Wild type | Diploid, 2459x2460 |
| DCY 4707 | a/$\alpha$ | T3A heterozygote | Diploid, 2459x4595 |
| DCY 4730 | a/$\alpha$ | T3A homozygote | Diploid, 4595x4596 |
| DCY 4824 | a/$\alpha$ | T3A heterozygote | Diploid, 2460x4596 |
| DCY 4963 | $\alpha$ | $\Delta lys2$ | 2460 *lys2::kan* transformant |
| DCY 4965 | $\alpha$ | $\Delta tyr7$ | 2460 *tyr7::kan* transformant |
| DCY 4970 | a | $\Delta tyr7$ | 2459 *tyr7::kan* transformant |
| DCY 4973 | $\alpha$ | T3A, $\Delta lys2$ | 4595 *lys2::kan* transformant |
| DCY 4979 | a | T3A, $\Delta tyr7$ | *hht1-hhf1::HYG<sup>R</sup>, hht2-T3A-HHF2(URA3), tyr7::kan* |
| DCY 4983 | a/$\alpha$ | Wild type | Diploid, 4963x4970 |
| DCY 4985 | a/$\alpha$ | T3A heterozygote | Diploid, 4963x4979 |
| DCY 4987 | a/$\alpha$ | T3A heterozygote | Diploid, 4970x4973 |
| DCY 4989 | a/$\alpha$ | T3A homozygote | Diploid, 4973x4979 |
| DCY 4751 | $\alpha$ | K4A | *hht1-hhf1::HYG<sup>R</sup>, hht2-K4A-HHF2(URA3)* |
| DCY 4752 | a | K4A | *hht1-hhf1::HYG<sup>R</sup>, hht2-K4A-HHF2(URA3)* |
| DCY 4753 | a | S10A | *hht1-hhf1::HYG<sup>R</sup>, hht2-S10A-HHF2(URA3)* |
| DCY 4754 | $\alpha$ | S10A | *hht1-hhf1::HYG<sup>R</sup>, hht2-S10A-HHF2(URA3)* |
| DCY 5043 | a/$\alpha$ | S10A heterozygote | Diploid, 2460x4753 |
| DCY 5045 | a/$\alpha$ | S10A heterozygote | Diploid, 2459x4754 |
| DCY 5047 | a/$\alpha$ | S10A homozygote | Diploid, 4753x4754 |
| DCY 5050 | a/$\alpha$ | K4A heterozygote | Diploid, 2459x4751 |
| DCY 5052 | a/$\alpha$ | K4A heterozygote | Diploid, 2460x4752 |
| DCY 5054 | a/$\alpha$ | K4A homozygote | Diploid, 4751x4752 |
| DCY 5058 | a | $\Delta mad2$ | *mad2::kan* |
| DCY 5061 | $\alpha$ | $\Delta mad2$ | *mad2::kan* |
| DCY 5064 | $\alpha$ | T3A, $\Delta mad2$ | *hht1-hhf1::HYG<sup>R</sup>, hht2-T3A-HHF2(URA3), mad2::kan* |
| DCY 5067 | a | T3A, $\Delta mad2$ | *hht1-hhf1::HYG<sup>R</sup>, hht2-T3A-HHF2(URA3), mad2::kan* |
| DCY 5074 | a/$\alpha$ | $\Delta mad2$ | Diploid, 5058x5061 |
| DCY 5077 | a/$\alpha$ | T3A, $\Delta mad2$ | Diploid, 5064x5067 |

### Yeast media

Three types of growth media were used in this study: YPD (Yeast extract, Peptone, Dextrose) medium, SD (Synthetic Defined) drop-out medium, and sporulation medium. YPD medium consisted of 10g/L Bacto-Yeast Extract, 20g/L Bacto-Peptone, 20g/L dextrose, and 20g/L agar. SD drop-out media consisted of 10g/L potassium acetate, 1g/L Yeast Extract, 0.5g/L dextrose, 20g/L agar, 6mg/L adenine, and 1.4g/L drop-out mix (1g L-adenine, 1g L-uracil, 2g L-tryptophan, 1g L-histidine, 1g L-arginine, 1g L-methionine, 3g L-tyrosine, 4g L-leucine, 4g L-isoleucine, 3g L-lysine, 2.5g L-phenylalanine, 5g L-glutamate, 5g L-aspartate, 7.5g L-valine, 10g L-threonine, and 20g L-serine, with the drop-out component excluded from the drop-out mix, i.e. SD-lys did not include the 3g L-lysine), with 0.5mL 4M NaOH added per liter of medium. Sporulation medium consisted of 10g/L potassium acetate, 1g/L Yeast Extract, 0.5g/L dextrose, 20g/L agar, and 6mg/L adenine.

### Yeast strain mating

Forced mating was employed to generate crosses of different yeast strains. To do this, two yeast cells of opposite mating type were placed next to each other on solid YPD medium using a dissection microscope. After two days of growth at 30°C, putative diploids were examined for the ability to produce mating pheromone to suppress haploid cell growth in halo assay tests.

### Mating-type (halo) assay

To determine mating type of non-diploid colonies, tester strains that arrest cell growth in the presence of the opposite mating factor were plated on solid YPD medium. Small amounts of the yeast isolate being tested were transferred to the test plate and grown at 30°C for two days. If the yeast isolate was of the opposite mating type to the tester, a halo where no tester cells could grow formed around the patch of yeast isolate. Diploid colonies do not form a halo on either the *a* or *α* test plates since they do not secrete mating factors.

### Diploid yeast sporulation

Yeast cells were grown in liquid YPD medium overnight at 30°C. The cells were then washed with ddH_2_O and plated on solid sporulation medium at room temperature. The cells were grown for three days on sporulation medium before sporulation analysis and tetrad dissection.

### Sporulation analysis

Sporulated cells were counted using a light microscope. The *Sporulation Efficiency* was calculated by categorizing cells as either spores or non-spores. Spores were defined as any cell that had attempted sporulation. These included cells that had completed the process of sporulation and had formed a mature ascus with either two, three or four spores inside the ascus. Also included were cells that had not completed ascus maturation but had either two, three or four spores within the round cell wall. Within each group, we also quantified *Successful Meiosis* (Table 2). *Successful Meiosis* was defined as the proportion of the spores that were tetrads with complete ascus maturation, i.e. as opposed to mature two, three or four spore asci, or immature cells that had begun but not completed sporulation.

**Table 2.**
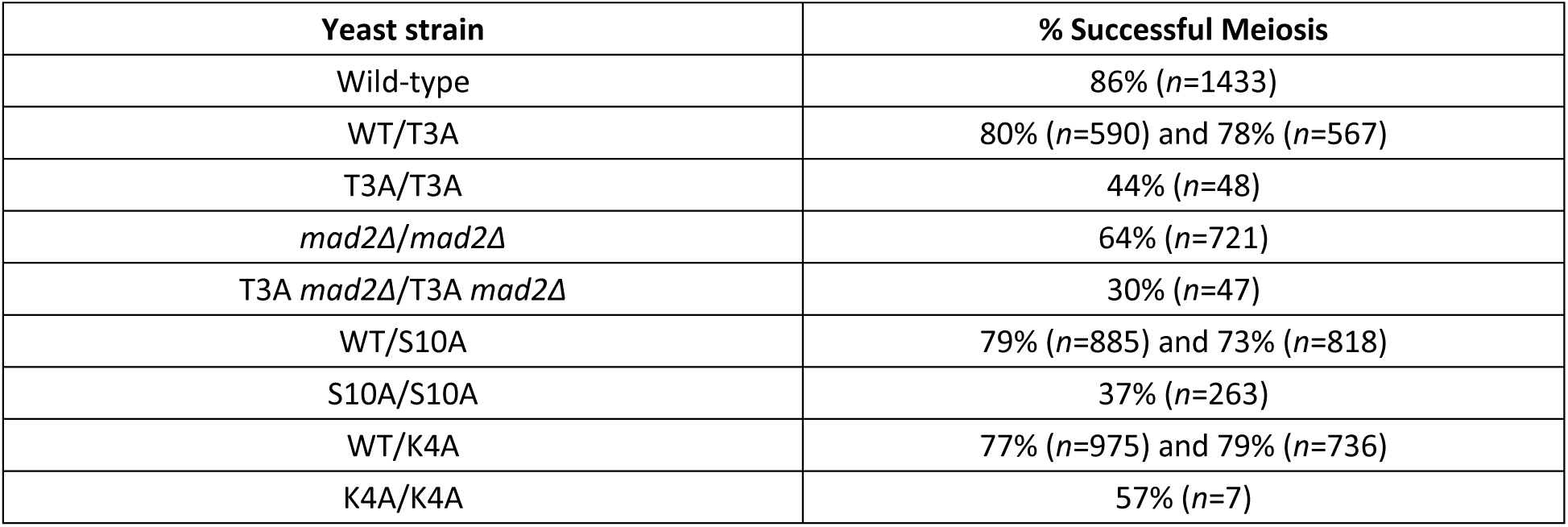
Quantification of successful meiosis. Total cells (*n*) attempting sporulation were categorized as having completed meiosis successfully (formation of mature tetrads) versus aberrantly (see Methods for full description of the aberrant category).

| Yeast strain | % Successful Meiosis |
| --- | --- |
| Wild-type | 86% (n=1433) |
| WT/T3A | 80% (n=590) and 78% (n=567) |
| T3A/T3A | 44% (n=48) |
| *mad2Δ/mad2Δ* | 64% (n=721) |
| T3A *mad2Δ*/T3A *mad2Δ* | 30% (n=47) |
| WT/S10A | 79% (n=885) and 73% (n=818) |
| S10A/S10A | 37% (n=263) |
| WT/K4A | 77% (n=975) and 79% (n=736) |
| K4A/K4A | 57% (n=7) |

### Tetrad analysis

Sporulated cells were incubated with 1 mg/mL lyticase for 12 minutes at 37°C before being plated on YPD medium. Tetrads were then dissected using a dissection microscope and grown for three days at 30°C. The resulting individual colonies were counted to determine spore viability.

### PCR and gel electrophoresis

PCR was done using 10 μL 10X PCR buffer (500mM KCl, 100mM Tris-HCl, pH 8.3 at 25°C), 1 μL dNTPs, 2.5 μL 3’ primer, 2.5 μL 5’ primer, 0.5 μL Taq DNA polymerase, 33 μL ultrapure water, and 0.5 μL target DNA. The primers were either *LYS2-kan* or *TYR7-kan* 5’ and 3’ primers. The PCR program used consisted of 30 cycles of a 30 second denaturation step at 94°C, 1 minute annealing step at 60°C, and 1 minute extension step at 72°C. The initial denaturation was 2 minutes at 95°C, and the final extension was 7 minutes at 72°C. Gel electrophoresis was used to analyze PCR products. For gel electrophoresis, a 0.8% agarose gel was run for 45 minutes in 1X TBE buffer (10.8g/L Tris, 5.5g/L boric acid, 4mL 500mM EDTA, pH 8.0).

### Yeast DNA transformation

5 mL liquid cultures in 1X adenine + YPD (4.95mL liquid YPD with 50μL 100X adenine) were grown overnight at 30°C, then diluted to OD_600_ ∼0.2 and grown for four hours. The cells were centrifuged, and the cell pellet was washed with ddH_2_O several times before being resuspended in 0.1 M LiAc. 12 μL of 10 mg/mL single-stranded salmon sperm (SSSS) DNA and 20 μL of PCR product were added to the cells, which were incubated at 30°C for 15 minutes. 600 μL LiAc/PEG (40 % PEG, mw ∼36,500, in 10 mM Tris-HCl, pH 8.0, with 1 mM EDTA and 0.1 M Lithium actetate) was added, and the cells were incubated again at 30°C for 30 minutes. 68 μL DMSO was added, and the cells were heat shocked at 42°C for 15 minutes. The cells were then centrifuged and resuspended in 500 μL YPD and grown for 1 hour at 30°C. The cells were then centrifuged and excess supernatant was discarded. The cell pellet was resuspended in the remaining supernatant and plated on YPD+G418 medium (YPD with 200mg/L G418 sulfate added) and grown at 30°C for two days.

### Replica plating

Velvet squares were used to make replicas of the colonies on one plate and transfer them to another plate. This was used to evaluate chromosome segregation by using SD-tyr and SD-lys plates where the cells with tyrosine and lysine auxotrophies, respectively, could not grow, allowing the segregation of these trackable markers to be followed in the daughter cells after meiosis and sporulation.

## Results

### Sporulation efficiency of yeast cells lacking histone H3 threonine 3 phosphorylation

To evaluate possible requirements of H3T3ph in meiosis, diploid yeast strains were made by crossing two wild-type strains to generate a wild-type homozygous diploid (DCY4706), crossing a wild-type and a T3A mutant to generate heterozygous T3A mutants (DCY4707 and DCY4824), and crossing two T3A strains to generate a homozygous T3A mutant (DCY4730) (Table 1). These diploids were plated on sporulation medium, and the sporulation efficiency was quantified by light microscopy after three days (Figure 1). Sporulation efficiency was 38.9% in the wild-type homozygotes, but was drastically reduced to only 1.9% in the homozygous T3A mutant where H3T3 cannot be phosphorylated. The two different strains heterozygous for the T3A substitution had sporulation efficiencies of 38.8% and 37.7%, very similar to the wild-type. The low sporulation frequency of the T3A homozygote suggests that the lack of H3T3ph negatively impacts the ability of the cells to initiate or complete meiosis, but the sporulation process in yeast cells can tolerate having only one-half of their histones with phosphorylatable H3T3.

**Figure 1.**
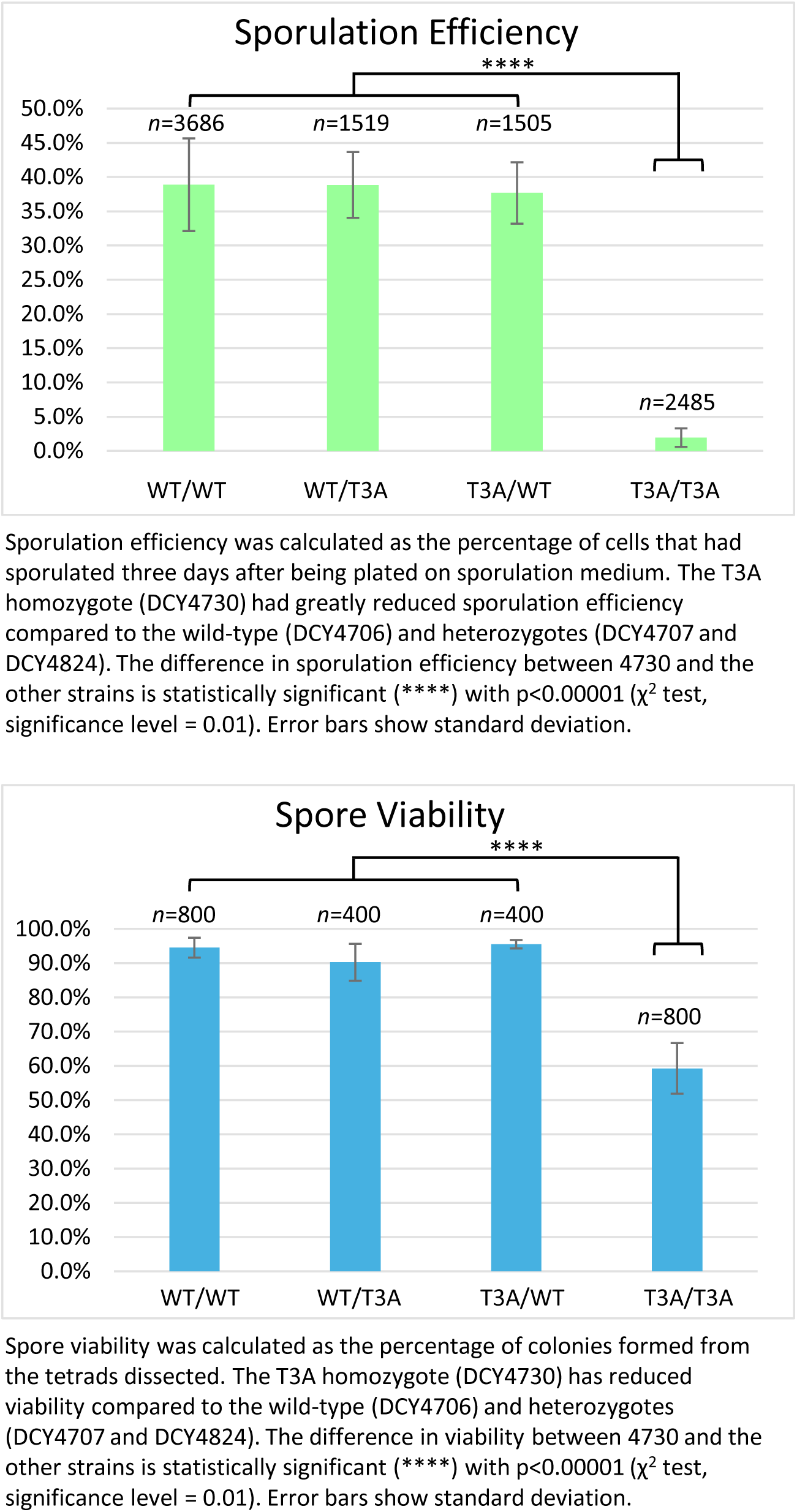
Histone H3 T3A Sporulation Efficiency and Spore Viability.

Next, we asked if the fidelity of meiosis was impacted in the T3A mutant (Table 2). Cells that attempted meiosis were categorized as having successfully completed meiosis if a mature four-spore ascus was produced (tetrad). Unsuccessful meiosis was defined as two-spore or three-spore asci that had completed sporulation or cells that had immature asci containing pro-spores (delayed or arrested part-way through meiosis). Based on these categories, out of 1433 wild-type cells attempting sporulation, 86% had completed meiosis successfully. The heterozygotes completed meiosis successfully 80% and 78% of the time. Of 48 homozygous T3A cells attempting sporulation, only 44% completed meiosis successfully. Therefore, not only was sporulation efficiency reduced in the T3A mutant, but successful completion of meiosis was perturbed.

### Spore viability in yeast cells lacking H3T3ph

Sporulation efficiency of the T3A homozygote was only 1.9% and of those only 44% produced mature tetrads. This indicates a critical role for this histone H3 residue in meiosis. We next asked if mature tetrads produced by T3A mutant cells were viable. Tetrads were dissected to separate each spore and allow them to grow into individual colonies. We dissected at least 100 tetrads for each strain and then quantified the number of spores that grew into viable colonies (Figure 1). This revealed 94.5% of wild-type spores were viable, while only 59.3% of the homozygous T3A spores were viable. The viability of the heterozygotes was similar to the wild-type, at 90.3% and 95.5%. We then quantified the frequency with which tetrads had all four spores giving rise to viable colonies, i.e. the percentage of viable tetrads. Tetrad viability was 87.5% for wild-type, 81% and 90% for the heterozygotes, but only 27.5% for the T3A mutant. Altogether, the T3A mutant had greatly reduced sporulation efficiency, a reduction in the frequency of sporulation resulting in the formation of mature four-spore asci, and significantly lower viability of the spores in those mature tetrads.

### H3 N-terminal tail mutations differentially affect spore viability

We next investigated if the effects the T3A mutation had on meiosis are specific to the threonine 3 residue or if mutation of other histone H3 tail residues have the same consequences. Homozygous and heterozygous histone H3 K4A and S10A mutants were generated, and their sporulation efficiency and spore viability were quantified as for the T3A mutants. We chose Lysine 4 because it is directly adjacent to threonine 3, however rather than being phosphorylated it can be methylated or acetylated. K4 methylation has been extensively studied and is known to be involved in regulation of gene expression pathways initiated during meiosis [27–31] as well as being associated with sites of meiotic recombination initiation in yeast [32, 33]. We chose S10 because, although it is farther away from T3, it is also subject to phosphorylation and plays known roles in chromosome compaction during both mitosis and meiosis [34–39].

Compared to the wild-type, the S10A mutant had reduced sporulation efficiency (10.1%) and reduced spore viability (75.0%) (Figure 2), but these effects on meiosis were not as severe as the T3A mutant. The frequency of successful meiosis was also reduced in the S10A mutant (Table 2). The K4A mutant had the lowest sporulation efficiency, 0.4% (Figure 3). It had such a low sporulation efficiency that tetrad dissection and analysis were impractical. These differences between T3A, S10A, and K4A show that each mutation has a different effect on meiosis (Statistical analysis is summarized in Figure 4). However, it is notable that all the mutations tested had deficits in sporulation efficiency and spore viability. Overall, the results suggest that H3T3ph has a distinct role in meiosis compared to H3 S10 phosphorylation and H3 K4 methylation/acetylation, indicating that the function of H3T3ph in meiosis is not synonymous with a general function of the H3 N-terminal tail.

**Figure 2.**
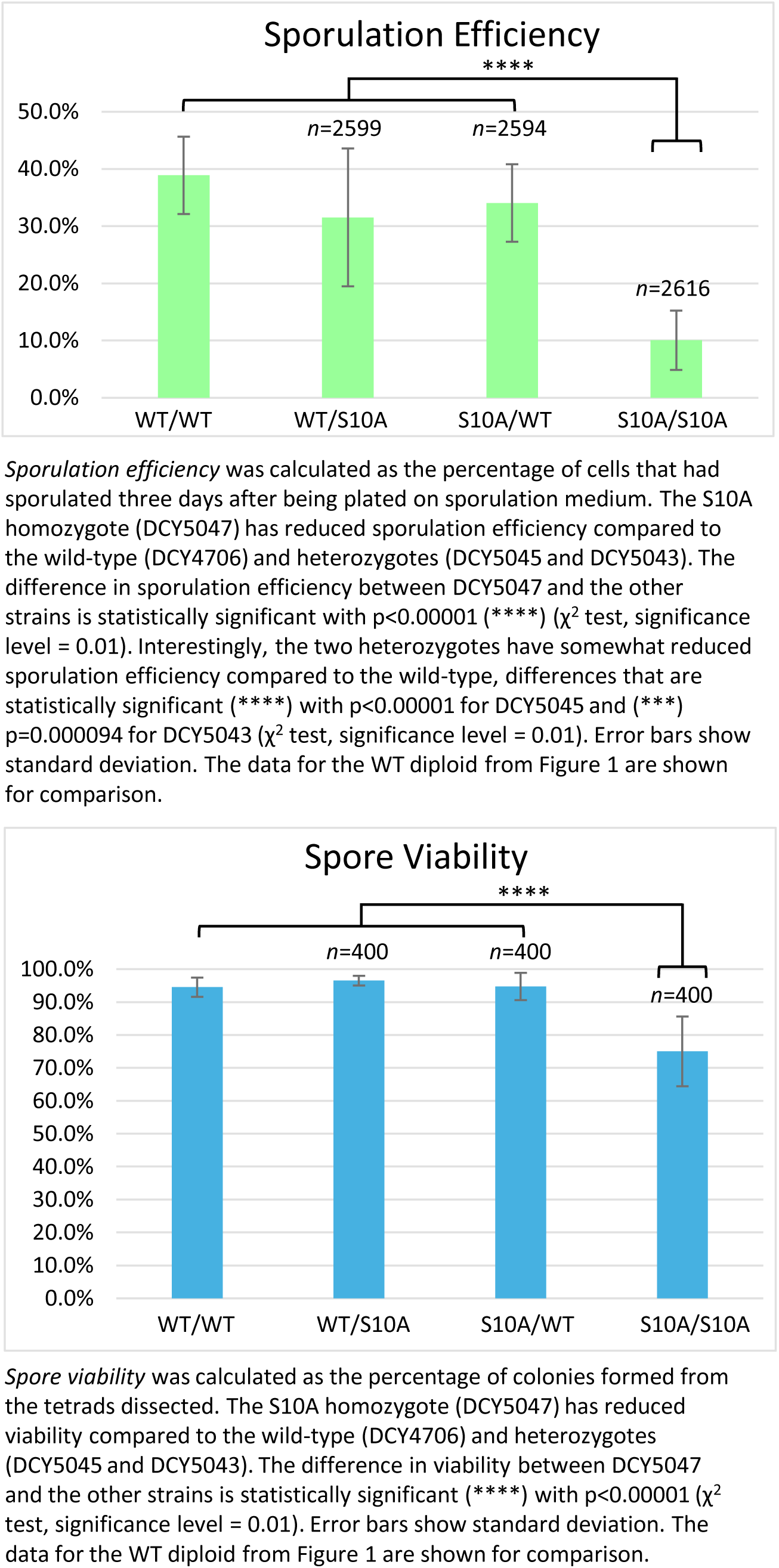
Histone H3 S10A Sporulation Efficiency and Spore Viability.

**Figure 3.**
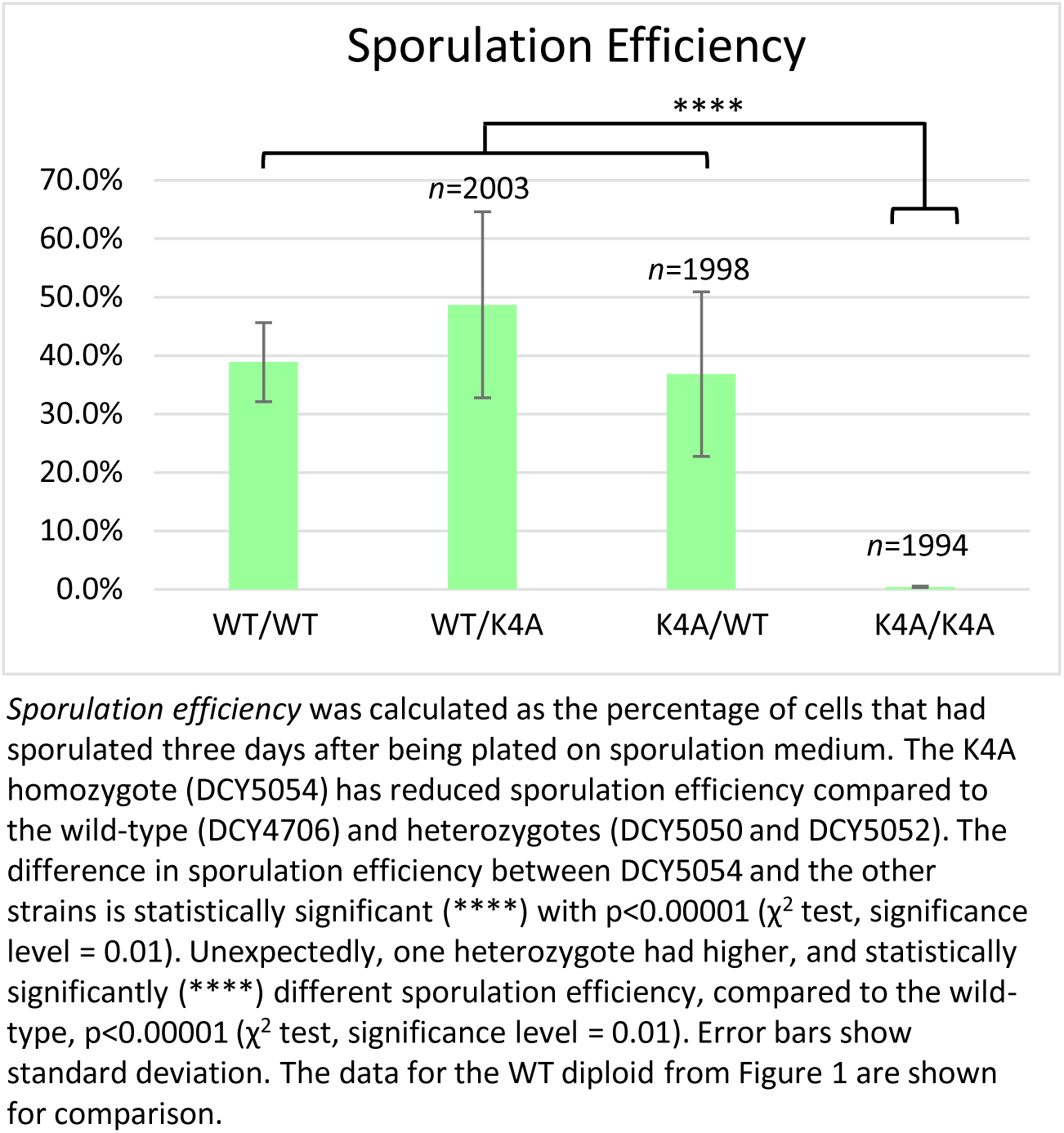
Histone H3 K4A Sporulation Efficiency.

**Figure 4.**
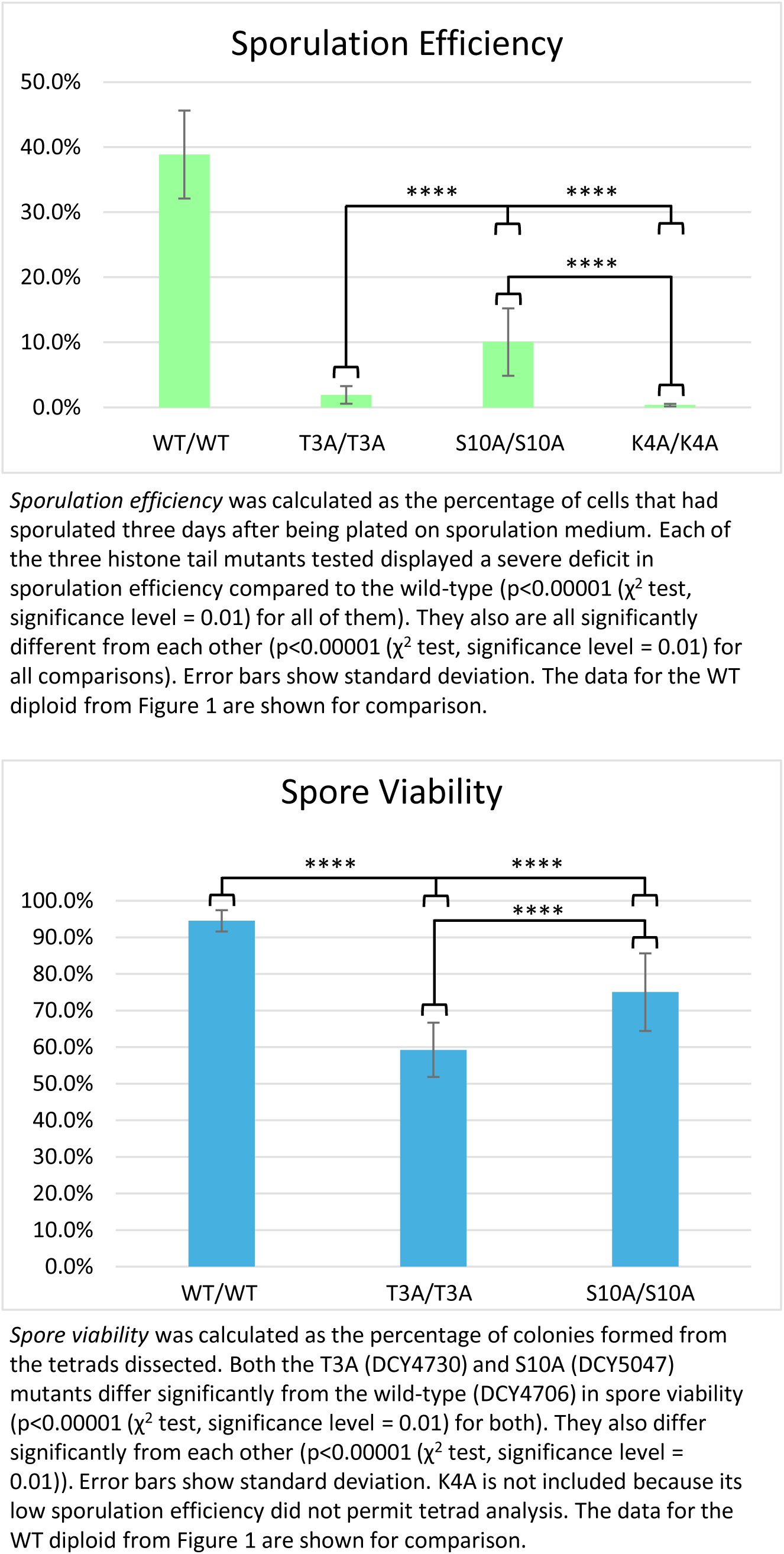
Comparison of Sporulation Efficiency and Spore Viability in the Histone H3 N-terminal tail mutants.

### H3T3 phosphorylation is not essential for proper recombination during meiosis

T3A homozygotes had very poor sporulation efficiency. Diploids that attempted sporulation produced mature asci at a reduced frequency and those that formed complete tetrads had low viability. These defects could be caused by aberrant recombination. To test this possibility, lysine and tyrosine auxotrophies were introduced into the haploid wild-type and T3A strains to serve as trackable genetic markers to assess recombination fidelity. Diploid strains were then isolated and tetrads dissected following sporulation. Based on the segregation patterns of the auxotrophic markers, aberrant recombination events were rare in all the strains examined. The wild-type had 99.5% proper segregation of the genetic makers, and the homozygous T3A mutant had 97.8% proper segregation (Figure 5). No significant difference was seen between the wild-type and T3A mutant. The heterozygotes also showed no significant difference at 99.5% and 98.3% proper segregation. Therefore, H3 T3 is not essential for the ability of *S. cerevisiae* cells to complete meiotic recombination successfully.

**Figure 5.**
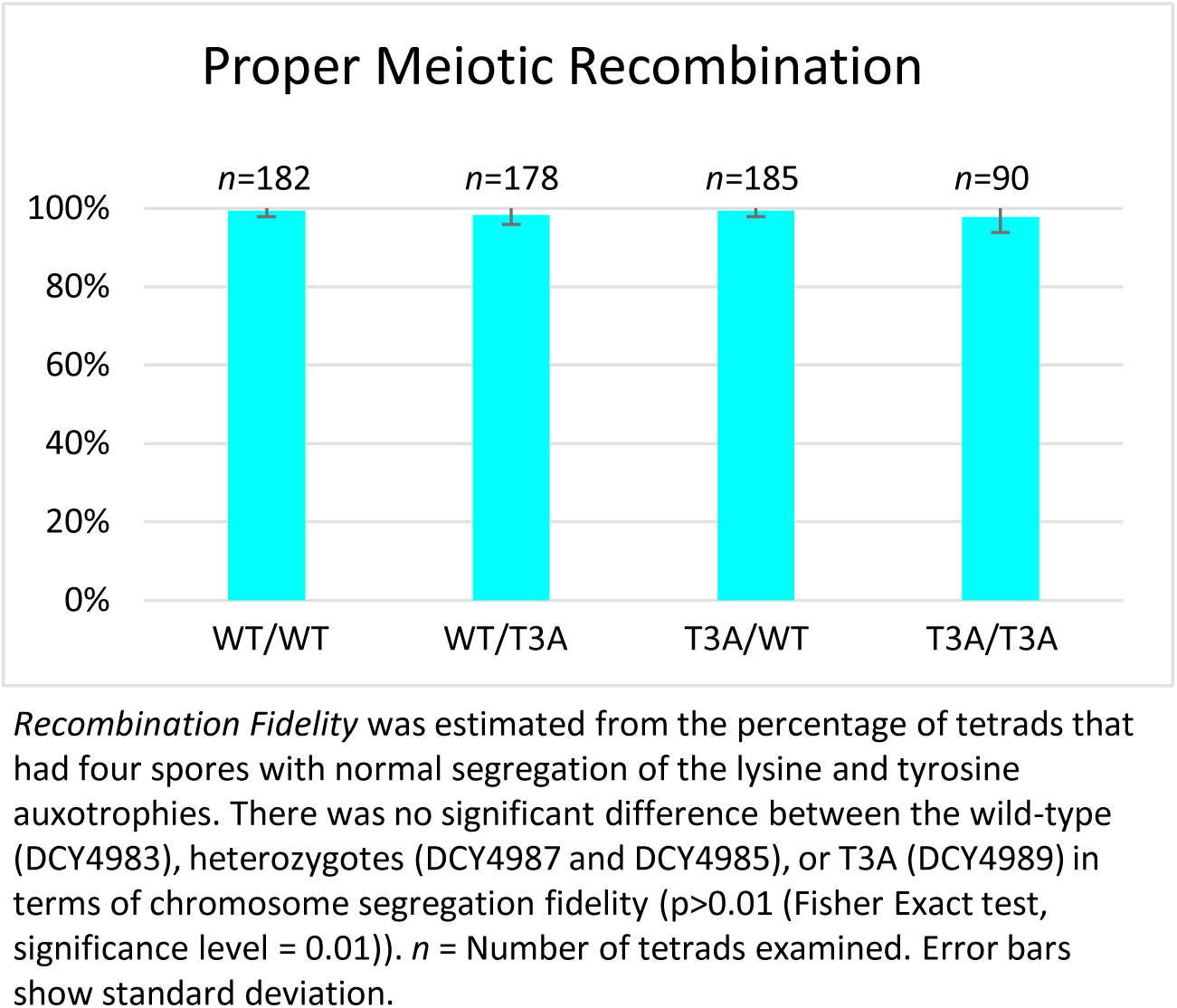
Analysis of Meiotic Chromosome Segregation Fidelity in the H3 T3A mutant.

### The spindle assembly checkpoint rescues spore viability in meiosis in the absence of H3 T3 phosphorylation

The T3A mutant had reduced spore viability in tetrads following meiosis, but this could not be explained by aberrant recombination. Dramatic phenotypes in the T3A mutant were reduced sporulation efficiency and formation of aberrant asci in cells that attempted sporulation. We reasoned that these phenotypes could be associated with activation of the spindle assembly checkpoint (SAC), because in mitosis H3T3ph is required to promote proper microtubule-kinetochore interactions. Lack of H3T3ph therefore activates the SAC in mitosis. The low sporulation efficiency and low spore viability in the T3A mutant might be due to improper kinetochore attachments, in which case the SAC might be triggered. The Mad2 protein is known to be required for the SAC in both mitosis and meiosis [26]. Therefore, we tested if sporulation efficiency and spore viability are compromised in T3A mutant cells lacking Mad2. A *mad2Δ* knockout mutation was crossed into the wild-type and the T3A mutant. We then quantified sporulation efficiency and spore viability for *mad2Δ* homozygotes and *mad2Δ* T3A homozygotes (Figure 6). The sporulation efficiency of the *mad2Δ* mutant was 35.7%, which is not significantly different from the wild-type (38.9%). The sporulation efficiency of the *mad2Δ* T3A double mutant was 2.4%, which is not significantly different from the T3A mutant (1.9%). Therefore, it does not seem that the SAC is restraining T3A mutants from progressing into or through the early stages of meiosis.

**Figure 6.**
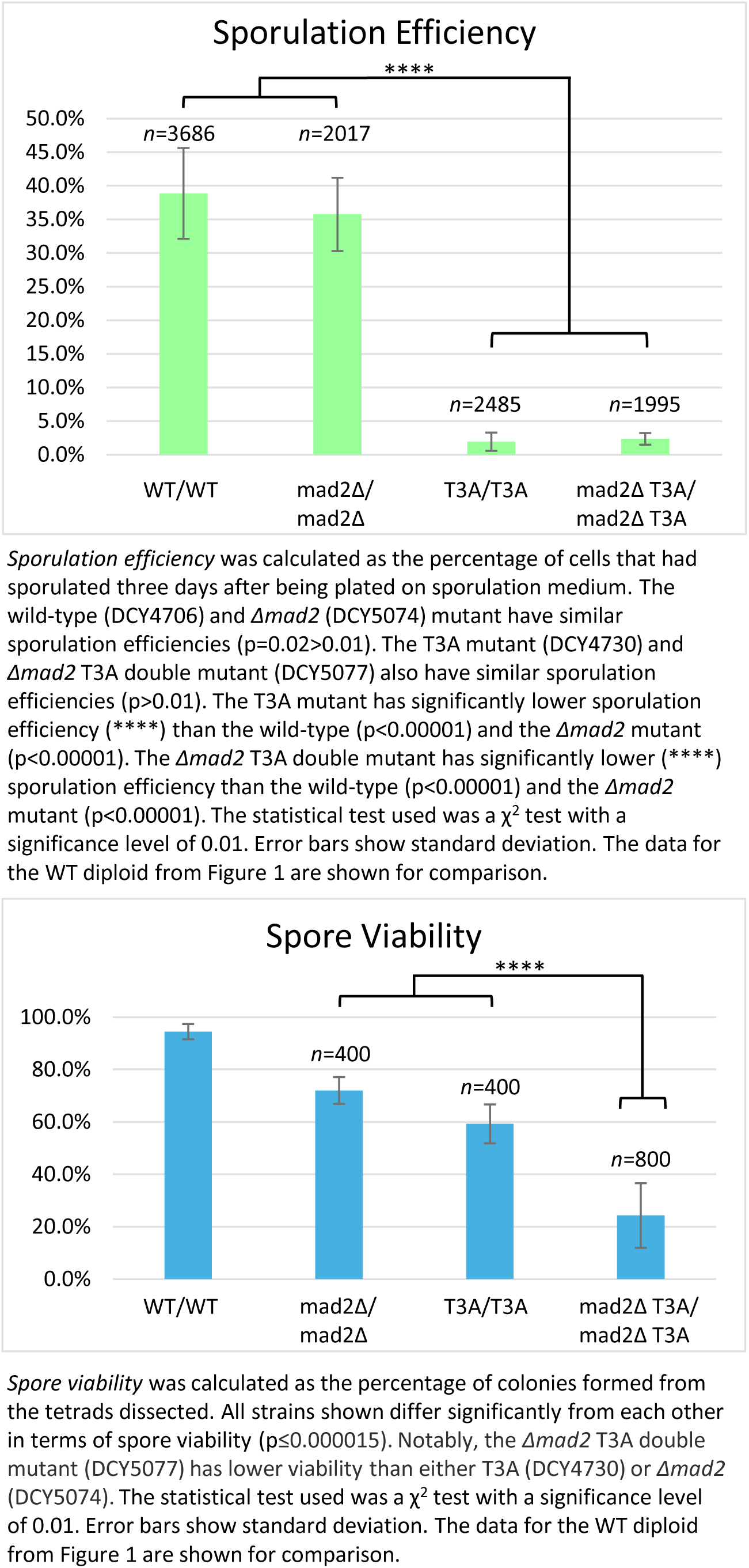
Histone H3 T3A Sporulation Efficiency and Spore Viability in combination with *MAD2* deletion.

The spore viability of the *mad2Δ* mutant was 72.0% compared to 94.5% in wild-type cells. This is consistent with the meiotic spindle checkpoint protecting cells from spore inviability. The spore viability of the *mad2Δ* T3A double mutant was only 24.3%, compared with 59.3% in the T3A mutant. The product of spore viabilities in the *mad2Δ* and T3A mutants (72% x 59.3%) is 42.7%. Therefore, the spore viability of the *mad2Δ* T3A double mutant (24.3%) is lower than would be expected if Mad2 and H3T3ph have unrelated functions in meiosis. This suggests that the SAC is involved in partially rescuing the meiotic defects in T3A cells.

### Spore viability patterns suggest the spindle assembly checkpoint rescues meiotic chromosome segregation errors in the absence of H3 T3 phosphorylation

The viability of spores in H3 T3A mutants partly depended on the spindle assembly checkpoint that prevents chromosome mis-segregation. Inviability of haploid yeast can be due to chromosome segregation errors because chromosome loss in haploid cells is lethal. The pattern of spore inviability after tetrad dissection can reveal chromosome segregation errors in meiosis I versus meiosis II. Mis-segregation of homologs in meiosis I results in two inviable spores, or four inviable spores if more than one chromosome mis-segregates. In contrast, mis-segregation of a sister chromatid in meiosis II results in three viable spores and one inviable spore. Analysis of spore viability patterns in H3 T3A mutants revealed that the largest increase was in the two inviable spore category (Figure 7). This was also the case for *mad2Δ* mutants. These patterns are most consistent with meiosis I single homolog segregation errors. In the T3A *mad2Δ* double mutant, four-spore inviability was most frequent, consistent with mis-segregation of multiple homologs in meiosis I.

**Figure 7.**
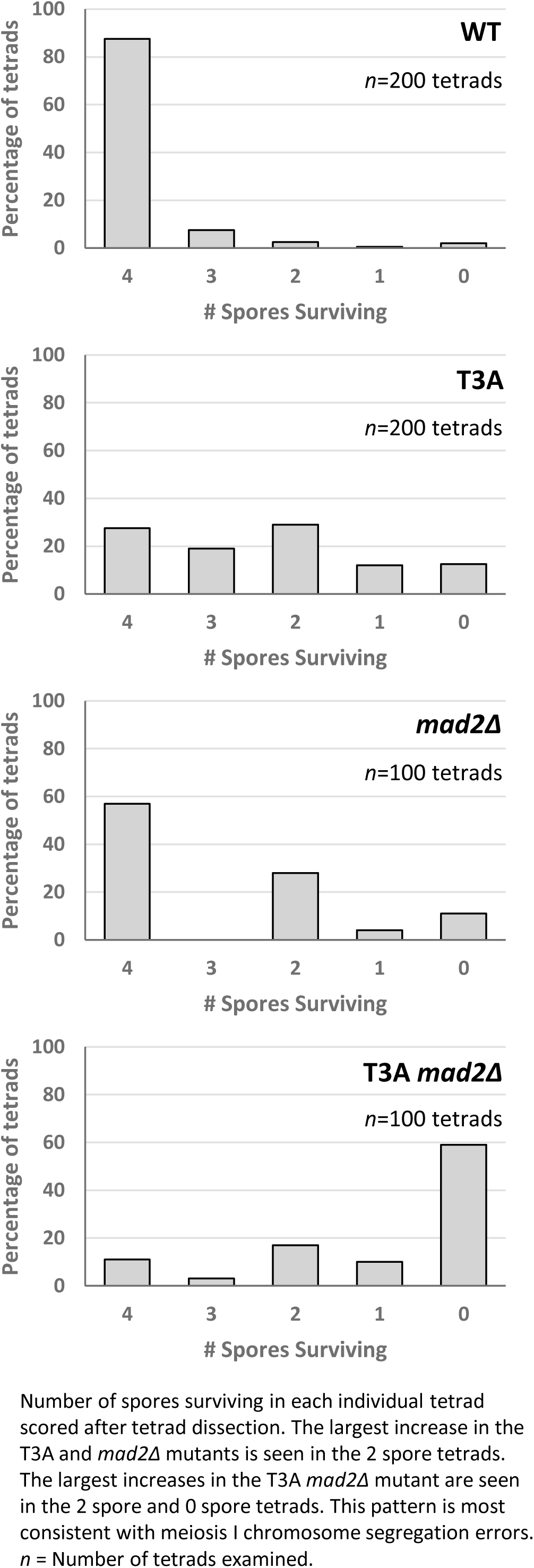
Tetrad Spore Analysis after Meiosis in H3 T3A mutants.

## Discussion

Phosphorylation of histone H3 at residue threonine 3 was recently detected in mitotic yeast cells and shown to be important for recruitment of Ipl1/Aurora B kinase to centromeres, as is the case in higher eukaryotes [15–17, 21]. Whether H3 T3 phosphorylation is important in meiosis has not been directly studied. Here, we investigated a possible requirement for H3T3ph during meiosis and have discovered that this residue plays a key role in meiotic progression and fidelity. We observed that T3A homozygous mutants had decreased sporulation efficiency, and cells that did attempt sporulation often produced aberrant spores or were delayed or arrested as immature spores. In cells that produced mature tetrads, spore viability relative to the wild type was significantly reduced. T3A heterozygotes behaved like wild-type cells, implying that these are recessive phenotypes. Assuming the heterozygotes express wild-type H3 and H3 T3A at approximately equal levels, this indicates that having reduced, but not eliminated, H3T3ph is tolerated in meiosis. However, a lack of H3T3ph is strikingly detrimental to the proper completion of meiosis, revealed by severe deficits in sporulation efficiency and in spore viability in the homozygous T3A mutant. Interestingly, inhibition of Haspin kinase (which targets H3 T3) in mice perturbed meiosis in similar ways to the T3A mutation in yeast. In mice, Haspin inhibition delayed the onset of meiosis I by delaying progression from prophase I at the step of nuclear envelope breakdown [23, 24]. Progression into anaphase I was also delayed, consistent with the observed defects in kinetochore-microtubule attachment [23]. Haspin inhibition also led to aneuploidy after meiosis in the mouse oocytes. This is consistent with our finding that yeast T3A mutants had reduced spore viability in mature tetrads.

We also investigated if the effects of the T3A substitution were specific to that mutation or if altering other residues within the histone H3 N-terminal tail have the same consequences. By assaying sporulation efficiency and spore viability in histone H3 K4A and S10A mutants, we were able to determine that the effects on meiosis are specific to T3A and are not general to other H3 tail mutations. Specifically, we observed that the T3A mutant has a higher sporulation efficiency than the K4A mutant, but a lower sporulation efficiency than the S10A mutant (Figure 4). We also observed that the T3A mutant has a lower spore viability than the S10A mutant (Figure 1 and 2). Therefore, there are significant differences between the effects of the T3A, K4A, and S10A mutations in meiosis. This suggests mechanistic differences between the roles of these residues. Notably, while all three mutants show different impacts, all three mutations are deleterious to the proper completion of meiosis. Interestingly, the S10A heterozygotes also had slightly reduced sporulation efficiency relative to the wild-type (Figure 2). The heterozygotes did not have as severe a defect as the homozygous S10A mutant, but the difference between the heterozygotes and the wild-type is still statistically significant. Although the reason for this difference is unclear, it suggests the possibility that sporulation is more sensitive to the level of modification of S10 than H3. Finally, it is important to note that a previous study did not observe decreased sporulation efficiency in yeast K4A or S10A strains [25]. In that case, a different strain background (SK1) was employed that has very high wild-type sporulation efficiency. One explanation for these apparent differences is that the strains used here (W303) have a reduced wild-type sporulation efficiency which likely results in a sensitized genetic background allowing additional requirements for meiosis to be revealed.

Existing evidence suggests what roles histone H3 S10 phosphorylation and K4 methylation/acetylation play in meiosis, versus T3 phosphorylation. S10 is phosphorylated in meiosis and is associated with chromosome compaction [34, 40]. K4 is a known regulator of gene expression and may contribute to the meiosis-specific program of gene expression needed to induce sporulation and assure the fidelity of meiosis [41–44]. On the other hand, the most clearly defined roles of H3 T3 are to promote biorientation of chromosomes on the spindle, and to delay the onset of anaphase when chromosomes cannot be segregated accurately [21, 22, 45–53].

Beyond these histone tail residues, other residues have been shown to be important for proper meiosis. For example, a histone H4 S1A mutant had ∼40% sporulation efficiency relative to wild-type cells [40], albeit a milder deficiency than H3 T3A (4.8% sporulation efficiency relative to wild-type cells). It is perhaps possible that H3 S10A and H4 S1A mutants had a less severe phenotype than T3A because both of those residues are linked to chromosome compaction [34, 54], arguing that T3A plays a distinct role.

We attempted to elucidate how the T3A mutation leads to decreased sporulation efficiency and decreased spore viability. Tracking the segregation of auxotrophic markers did not reveal evidence of aberrant recombination in T3A mutants (Figure 5). We then wondered if the spindle checkpoint is activated in the T3A mutant: The reduced sporulation efficiency could be explained by the SAC delaying meiotic progression. However, sporulation efficiency was not increased in *mad2Δ* T3A double mutants, as would have been predicted. Nevertheless, we also observed that spore viability in *mad2Δ* T3A mutants (24.3%) is much lower than either the T3A mutant (59.3%) or the *mad2Δ* mutant (72.0%) (Figure 6). Therefore, the spindle assembly checkpoint appears to function in suppressing the spore viability deficit seen in T3A cells. Because the spindle assembly checkpoint is activated when there are improper spindle attachments [26], and because the purpose of H3T3ph in mitosis is to correct improper kinetochore-microtubule interactions [23], the data suggest the main role of H3T3ph in meiosis is likewise to promote error correction mechanisms to ensure chromosome biorientation. Interestingly, the spore viability patterns of H3 T3A tetrads and H3 T3A *mad2Δ* double mutant tetrads (Figure 7) were most consistent with homolog chromosome segregation errors in meiosis I. Investigating how the SAC in conjunction with H3T3ph modulate meiosis and how the spindle assembly checkpoint can rescue the inviability of T3A spores will be interesting future directions.

## Acknowledgements

This work was supported by NIH/NIGMS GM130858.

## References

1. Malik, S.B., et al., An expanded inventory of conserved meiotic genes provides evidence for sex in Trichomonas vaginalis. PLoS One, 2007. 3(8): p. e2879.

2. Neiman, A.M., Sporulation in the budding yeast Saccharomyces cerevisiae. Genetics, 2011. 189(3): p. 737–65.

3. Peterson, C.L. and M.A. Laniel, *Histones and histone modifications.* Curr Biol, 2004. **14**(14): p. R546–51.

4. Marino-Ramirez, L., et al., Histone structure and nucleosome stability. Expert Rev Proteomics, 2005. 2(5): p. 719–29.

5. Mersfelder, E.L. and M.R. Parthun, The tale beyond the tail: histone core domain modifications and the regulation of chromatin structure. Nucleic Acids Res, 2006. 34(9): p. 2653–62.

6. Gu, L., Q. Wang, and Q.Y. Sun, Histone modifications during mammalian oocyte maturation: dynamics, regulation and functions. Cell Cycle, 2010. 9(10): p. 1942–50.

7. Xu, D., et al., Covalent modifications of histones during mitosis and meiosis. Cell Cycle, 2009. 8(22): p. 3688–94.

8. Wang, L., et al., The histone codes for meiosis. Reproduction, 2017. 154(3): p. R65–R79.

9. Gao, J. and M.P. Colaiacovo, Zipping and Unzipping: Protein Modifications Regulating Synaptonemal Complex Dynamics. Trends Genet, 2018. 34(3): p. 232–245.

10. Tolsma, T.O. and J.C. Hansen, Post-translational modifications and chromatin dynamics. Essays Biochem, 2019. 63(1): p. 89–96.

11. Nie, H., et al., Roles of histone post-translational modifications in meiosisdagger. Biol Reprod, 2024. 110(4): p. 648–659.

12. Rossetto, D., N. Avvakumov, and J. Cote, Histone phosphorylation: a chromatin modification involved in diverse nuclear events. Epigenetics, 2012. 7(10): p. 1098–108.

13. Sundararajan, S., et al., Methylated histones on mitotic chromosomes promote topoisomerase IIalpha function for high fidelity chromosome segregation. iScience, 2023. 26(5): p. 106743.

14. Quadri, R., S. Sertic, and M. Muzi-Falconi, Roles and regulation of Haspin kinase and its impact on carcinogenesis. Cell Signal, 2022. 93: p. 110303.

15. Dai, J. and J.M. Higgins, Haspin: a mitotic histone kinase required for metaphase chromosome alignment. Cell Cycle, 2005. 4(5): p. 665–8.

16. Dai, J., et al., The kinase haspin is required for mitotic histone H3 Thr 3 phosphorylation and normal metaphase chromosome alignment. Genes Dev, 2005. 19(4): p. 472–88.

17. Yamagishi, Y., et al., Two histone marks establish the inner centromere and chromosome bi-orientation. Science, 2010. 330(6001): p. 239–43.

18. Sawicka, A. and C. Seiser, Histone H3 phosphorylation - a versatile chromatin modification for different occasions. Biochimie, 2012. 94(11): p. 2193–201.

19. Wang, F., et al., Histone H3 Thr-3 phosphorylation by Haspin positions Aurora B at centromeres in mitosis. Science, 2010. 330(6001): p. 231–5.

20. Kelly, A.E., et al., Survivin reads phosphorylated histone H3 threonine 3 to activate the mitotic kinase Aurora B. Science, 2010. 330(6001): p. 235–9.

21. Edgerton, H., et al., A noncatalytic function of the topoisomerase II CTD in Aurora B recruitment to inner centromeres during mitosis. J Cell Biol, 2016. 213(6): p. 651–64.

22. Yoshida, M.M., et al., SUMOylation of DNA topoisomerase IIalpha regulates histone H3 kinase Haspin and H3 phosphorylation in mitosis. J Cell Biol, 2016. 213(6): p. 665–78.

23. Nguyen, A.L., et al., Phosphorylation of threonine 3 on histone H3 by haspin kinase is required for meiosis I in mouse oocytes. J Cell Sci, 2014. 127(Pt 23): p. 5066–78.

24. Wang, Q., et al., H3 Thr3 phosphorylation is crucial for meiotic resumption and anaphase onset in oocyte meiosis. Cell Cycle, 2016. 15(2): p. 213–24.

25. Govin, J., et al., Systematic screen reveals new functional dynamics of histones H3 and H4 during gametogenesis. Genes Dev, 2010. 24(16): p. 1772–86.

26. Tsuchiya, D., C. Gonzalez, and S. Lacefield, The spindle checkpoint protein Mad2 regulates APC/C activity during prometaphase and metaphase of meiosis I in Saccharomyces cerevisiae. Mol Biol Cell, 2011. 22(16): p. 2848–61.

27. Burgess, S.M., M. Ajimura, and N. Kleckner, GCN5-dependent histone H3 acetylation and RPD3-dependent histone H4 deacetylation have distinct, opposing effects on IME2 transcription, during meiosis and during vegetative growth, in budding yeast. Proc Natl Acad Sci U S A, 1999. 96(12): p. 6835–40.

28. Choy, J.S., et al., Yng2p-dependent NuA4 histone H4 acetylation activity is required for mitotic and meiotic progression. J Biol Chem, 2001. 276(47): p. 43653–62.

29. Nislow, C., E. Ray, and L. Pillus, SET1, a yeast member of the trithorax family, functions in transcriptional silencing and diverse cellular processes. Mol Biol Cell, 1997. 8(12): p. 2421–36.

30. Sollier, J., et al., Set1 is required for meiotic S-phase onset, double-strand break formation and middle gene expression. EMBO J, 2004. 23(9): p. 1957–67.

31. Sarmento, O.F., et al., Dynamic alterations of specific histone modifications during early murine development. J Cell Sci, 2004. 117(Pt 19): p. 4449–59.

32. Borde, V., et al., Histone H3 lysine 4 trimethylation marks meiotic recombination initiation sites. EMBO J, 2009. 28(2): p. 99–111.

33. Buard, J., et al., Distinct histone modifications define initiation and repair of meiotic recombination in the mouse. EMBO J, 2009. 28(17): p. 2616–24.

34. Hendzel, M.J., et al., Mitosis-specific phosphorylation of histone H3 initiates primarily within pericentromeric heterochromatin during G2 and spreads in an ordered fashion coincident with mitotic chromosome condensation. Chromosoma, 1997. 106(6): p. 348–60.

35. Wei, Y., et al., Phosphorylation of histone H3 at serine 10 is correlated with chromosome condensation during mitosis and meiosis in Tetrahymena. Proc Natl Acad Sci U S A, 1998. 95(13): p. 7480–4.

36. Wei, Y., et al., Phosphorylation of histone H3 is required for proper chromosome condensation and segregation. Cell, 1999. 97(1): p. 99–109.

37. Hsu, J.Y., et al., Mitotic phosphorylation of histone H3 is governed by Ipl1/aurora kinase and Glc7/PP1 phosphatase in budding yeast and nematodes. Cell, 2000. 102(3): p. 279–91.

38. Wang, Q., et al., Histone phosphorylation and pericentromeric histone modifications in oocyte meiosis. Cell Cycle, 2006. 5(17): p. 1974–82.

39. Wang, Q., et al., The spatial relationship between heterochromatin protein 1 alpha and histone modifications during mouse oocyte meiosis. Cell Cycle, 2008. 7(4): p. 513–20.

40. Krishnamoorthy, T., et al., Phosphorylation of histone H4 Ser1 regulates sporulation in yeast and is conserved in fly and mouse spermatogenesis. Genes Dev, 2006. 20(18): p. 2580–92.

41. Primig, M., et al., The core meiotic transcriptome in budding yeasts. Nat Genet, 2000. 26(4): p. 415–23.

42. Mitchell, A.P., Control of meiotic gene expression in Saccharomyces cerevisiae. Microbiol Rev, 1994. 58(1): p. 56–70.

43. Chu, S. and I. Herskowitz, Gametogenesis in yeast is regulated by a transcriptional cascade dependent on Ndt80. Mol Cell, 1998. 1(5): p. 685–96.

44. Chu, S., et al., The transcriptional program of sporulation in budding yeast. Science, 1998. 282(5389): p. 699–705.

45. Berenguer, I., et al., Haspin participates in AURKB recruitment to centromeres and contributes to chromosome congression in male mouse meiosis. J Cell Sci, 2022. 135(13).

46. Cairo, G., et al., Distinct Aurora B pools at the inner centromere and kinetochore have different contributions to meiotic and mitotic chromosome segregation. Mol Biol Cell, 2023. 34(5): p. ar43.

47. Cairo, G. and S. Lacefield, Establishing correct kinetochore-microtubule attachments in mitosis and meiosis. Essays Biochem, 2020. 64(2): p. 277–287.

48. Hadders, M.A., et al., Untangling the contribution of Haspin and Bub1 to Aurora B function during mitosis. J Cell Biol, 2020. 219(3).

49. Johansson, M., Y. Azuma, and D.J. Clarke, Role of Aurora B and Haspin kinases in the metaphase Topoisomerase II checkpoint. Cell Cycle, 2021. 20(4): p. 345–352.

50. Kurihara, D., et al., Identification and characterization of plant Haspin kinase as a histone H3 threonine kinase. BMC Plant Biol, 2011. 11: p. 73.

51. Wang, F., et al., Haspin inhibitors reveal centromeric functions of Aurora B in chromosome segregation. J Cell Biol, 2012. 199(2): p. 251–68.

52. Pandey, N., et al., Topoisomerase II SUMOylation activates a metaphase checkpoint via Haspin and Aurora B kinases. J Cell Biol, 2020. 219(1).

53. Clarke, D.J. and Y. Azuma, Non-Catalytic Roles of the Topoisomerase IIalpha C-Terminal Domain. Int J Mol Sci, 2017. 18(11).

54. Barber, C.M., et al., The enhancement of histone H4 and H2A serine 1 phosphorylation during mitosis and S-phase is evolutionarily conserved. Chromosoma, 2004. 112(7): p. 360–71.

